# ViroSeek: a viral detection pipeline for second-generation sequencing

**DOI:** 10.64898/2026.03.04.706323

**Authors:** Audric Berger, Margaux J.M. Lefebvre, Jacques Dainat, Davy Jiolle, Isabelle Moltini-Conclois, Loic Talignani, Emilio Mastriani, Sylvie Cornelie, Nicolas Berthet, Christophe Paupy

## Abstract

Arbovirus emergences represent a rising public health issue and are exacerbated by climate change and globalization. Virome analysis has become a key approach for monitoring and managing infectious diseases, yet existing tools often remain technically complex and inaccessible to non-specialists. In this context, we present ViroSeek, a reproducible and accessible bioinformatics pipeline specifically designed for the taxonomic analysis of second-generation sequencing data from target-enriched libraries. ViroSeek performs a series of automated steps: quality control, trimming, host and bacterial sequence removal, assembly, taxonomic assignment, read remapping for quantification, and PCR duplicate removal. The whole process is designed to produce a clear, usable viral taxonomy table that is suitable for diversity studies. ViroSeek was empirically validated on enriched control samples containing a known panel of viruses. All the expected viruses were correctly detected. Bacterial and host contaminant sequences were effectively removed. The pipeline is freely available and fully documented, supporting its adoption and adaptation by the research community.

## Introduction

Viruses are a major global public health issue, particularly because of arbovirosis and zoonotic viruses (I. F. de Almeida et al. 2023; Dong and Soong 2021). Climate change, growing urbanization or population movements contribute to increasing the risks of viral emergence and transmission (Baker et al. 2022). In this context, virome analysis, which aims to study the viral diversity present in different environments or hosts, is now a major challenge in microbial ecology, human health and environmental monitoring (Quick et al. 2016; J. P. de Almeida et al. 2021; I. F. de Almeida et al. 2023).

Traditional virus characterization primarily depends on laboratory-based methods such as cell culture (Grandien and Svedmyr 1983). While effective, these approaches are time-intensive and generally restricted to the identification of individual viruses. In contrast, next-generation sequencing (NGS) has emerged as a high-throughput, cost-effective alternative for large-scale viral detection. Therefore, numerous bioinformatics tools have been developed to detect, classify and annotate viral sequences from metagenomic data (Kaiser et al. 2025).

However, despite their abundance, most of these pipelines have significant limitations. Some are too specialized (*i.e*, Metaphage or poreCov) (Pandolfo et al. 2022; Brandt et al. 2021), others are poorly maintained, difficult to install due to obsolete dependencies, or contain errors that prevent proper execution of the scripts. For example, a study comparing twenty virome pipelines for third-generation sequencing data revealed that only five of them were usable, the others being either unsuitable or impossible to operate (Kaiser et al. 2025).

Furthermore, several existing pipelines and tools are specifically designed for the detection of viruses infecting bacteria and archaea (Kieft et al. 2020; Guo et al. 2021; Ren et al. 2017), making them unsuitable for the identification of arboviruses. In this study, we focus on the analysis of viromes derived from second-generation (Illumina) sequencing data from target-enriched libraries. The pipeline is designed to handle both RNA and DNA viruses, with a particular emphasis on eukaryotic viruses, including arboviruses of public health concern.

Pipelines such as Taxprofiler (Stamouli et al. 2023), MetaDenovo (Lundin et al. 2025), PIMGAVir (Mastriani et al. 2022), MicroSeek (Pérot et al. 2022), or VirusTaxo (Raju et al. 2022) were initially considered. However, their integration proved challenging due to installation issues and output files were found to be unsuitable for some downstream analyses. In the case of MetaDenovo, assembly is performed across all samples collectively. However, retaining assemblies on a per-sample basis facilitates downstream analyses such as phylogenetic placement and genetic diversity assessment. Furthermore, MetaDenovo performs taxonomic assignment using *blastp*, which requires prior ORF prediction and may introduce biases in viral sequences (Finkel et al. 2018). Although *blastn* avoids this step, protein level searches provide higher sensitivity for divergent viruses (Pearson 2014). Therefore, *BLASTx, which* evaluates all six reading frames without explicit ORF calling, offers a more robust alternative (Buchfink et al. 2021). This approach is implemented when *Diamond* is selected as the taxonomic classifier in Taxprofiler. However, raw reads are not assembled at any stage of the workflow, which does not enable the phylogenetic placement of sequences assigned to specific species. To address this limitation, we adopted an assembly-based strategy, enabling the reconstruction of longer viral sequences and facilitating downstream analyses of genetic diversity. While VirusTaxo provides taxonomic classification of viral sequences, it does not include an assembly step and requires pre-assembled contigs as input, limiting its standalone utility in a complete pipeline. Finally, other available pipelines also present accessibility limitations. For instance, PIMGAVir presented challenges including high RAM requirements and restrictive access rights, while MicroSeek lacked publicly available code or workflow documentation.

Although flexible workflows such as MetaDenovo or ViralMetagenome provide comprehensive solutions for the analysis of diverse viral communities, they are designed to accommodate a broad range of datasets and experimental designs. Consequently, they offer extensive flexibility and multiple configurable steps in this objective. ViroSeek was developed for a more specific application: the analysis of short-read target-enriched libraries generated for the monitoring and characterization of eukaryotic viruses.

Rather than integrating multiple interchangeable tools, the design of Viroseek relies on two main methodological choices. An assembly-based strategy to generate sample specific viral contigs and a taxonomic assignment using protein-level similarity searches. Relying on a single optimized combination of methods to provide consistency and interpretable results. This approach remains accessible to non-expert users and devoid of unnecessary options while maintaining sensitivity for divergent eukaryotic viruses. To validate the pipeline, we analyzed four paired-end metagenomic libraries with known viral compositions, and one simulated dataset containing viral sequences with bacterial contaminants. The pipeline is available in a public GitHub repository (MargauxLefebvre/ViroSeek) and is thoroughly documented to facilitate installation, usage and potential modification by the community.

## Materials and methods

### Description of the pipeline

ViroSeek is a pipeline designed for the analysis of target-enriched libraries (sequencing libraries generated after an experimental enrichment step for predefined targets), with specific optimization for managing high PCR duplicate rates and performing per-sample assembly (Figure 1). The primary inputs are paired or single-end FASTQ files generated by NGS. The pipeline can be executed using a simple command such as: *nextflow run MargauxLefebvre/ViroSeek -r v0.0.2 --input ‘/path/to/samples.csv’*, with all parameters set to their default values.

**Figure 1:**
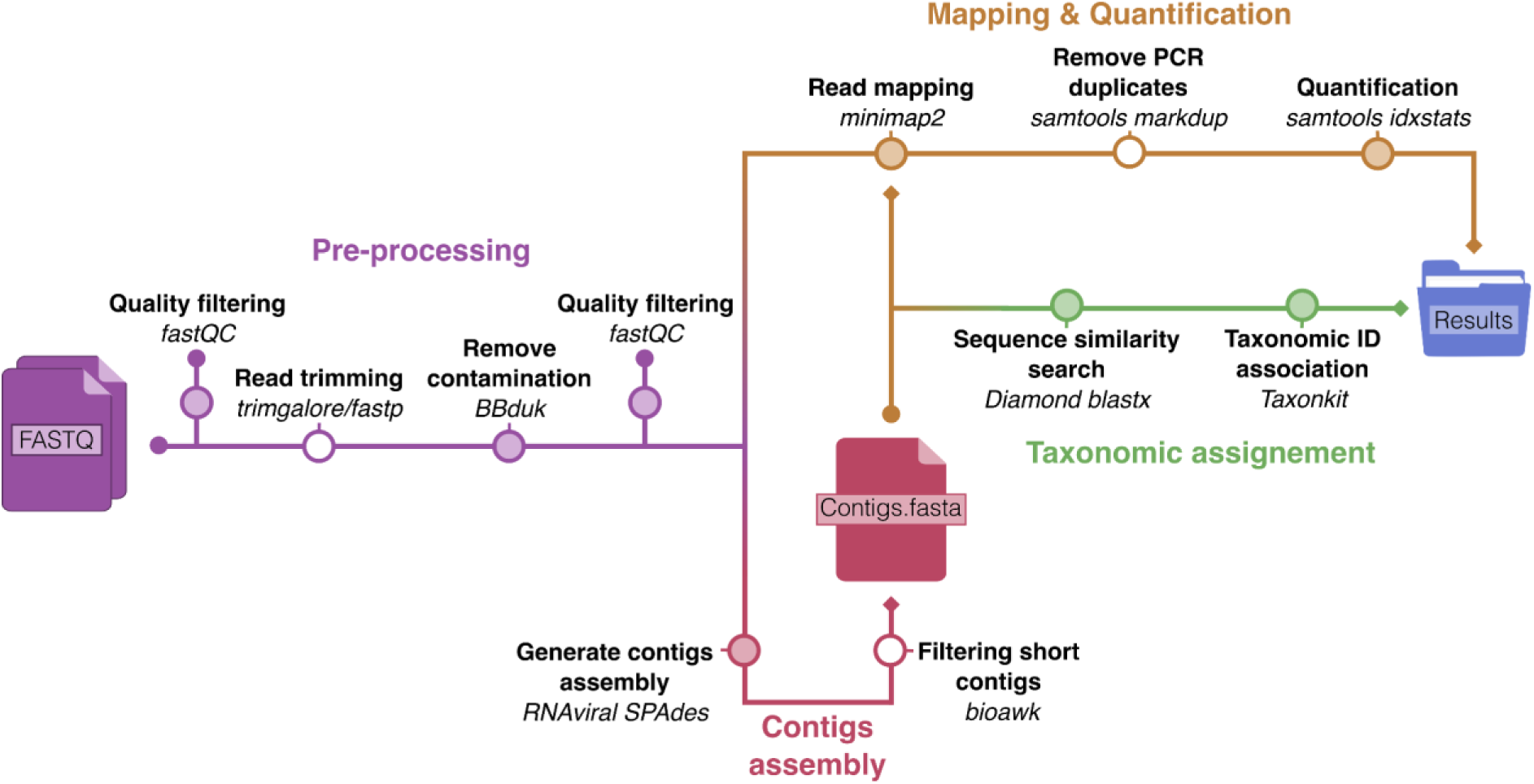
Simplified workflow of the ViroSeek pipeline. The pipeline comprises four main stages for the processing and analysis of target-enriched libraries: pre-processing, assembly construction, taxonomic assignment, and read mapping for relative quantification. For each step (shown in bold), the corresponding software tool is indicated in italics. The steps marked with a white circle are optional and can be skipped. The version used for each tool of the pipeline is described in Supplementary Table 1.

Raw sequencing reads from the paired-end or single-end FASTQ files are pre-processed using *Trim Galore v0.6.10* (Krueger et al. 2023) or *fastp* v0.23.4 (Chen et al. 2018; Chen 2025) to remove adapter sequences and low-quality bases. Subsequently, potential contamination is filtered out using *BBduk v39.18* (Bushnell 2026). By default, ViroSeek removes non-viral ribosomal RNA (16S/18S and 23S/28S) employing the SILVA rRNA database as a reference (Quast et al. 2013, release 138). However, users may substitute this database with an alternative reference better suited to their specific application, such as a host genome. Quality control is performed after each filtering step using *FastQC v0.12.1* (Andrews et al. 2023).

For each sample, de novo assembly is carried out using *SPAdes v4.2.0* (Prjibelski et al. 2020) with the *--rnaviral* option, which is specifically designed for assembling viral RNA-seq data sets. The assembly can be filtered to retain only contigs exceeding a specified minimum length specified by the user, thereby reducing incorrect taxonomic assignments associated with shorter contigs.

Taxonomic assignment of assembled contigs is performed using *Diamond v2.1.13* (Buchfink et al. 2021) in *BLASTx* mode, which aligns nucleotide contigs against a protein database. By default, ViroSeek uses a database derived from the NCBI reference sequence non-redundant protein database (Pruitt et al. 2005), restricted to viral sequences. However, users may specify an alternative database generated with the *makedb* utility from *Diamond v2.1.13.* The *--range-culling* option is enabled to allow *Diamond* to merge multiple high-scoring segment pairs from the same subject sequence, resulting in a more accurate estimation of total query coverage. The alignment is run in frameshift-aware mode (*-F 15*) to account for potential indels or sequencing errors that might cause frameshifts in the RNA viral genome. Moreover, user-defined thresholds can be specified for the maximum *e-value* of alignment hits, the minimum percent identity, and the minimum query coverage. In addition, the user can also define the sensitivity mode of *Diamond* (*faster*, *fast*, *mid-sensitive*, *sensitive*, *more-sensitive*, *very-sensitive* or *ultra-sensitive*). Assignment is followed by classification with *Taxonkit v0.9.0* (Shen and Ren 2021). To estimate the relative viral abundance (Roux et al. 2017; Kane et al. 2024), cleaned reads are mapped to the assembled contigs using *Minimap 2 v2.29* (Li 2018) and PCR duplicates are removed using the *markdup* function from *Samtools v1.21* (Danecek et al. 2021). However, this *Samtools* function relies exclusively on read position and orientation, which may result in the erroneous classification of genuine signals as duplicates by chance (Danecek et al. 2021). To mitigate this potential bias, users may bypass the deduplication step by specifying the -*-skip_dedup* flag.

All essential tools required by the pipeline are encapsulated within a *Singularity* (Kurtzer et al. 2017) or *Docker* (Merkel 2014) container to ensure full reproducibility across computing environments. The workflow is automatically managed by Nextflow *v24.10.4* (Di Tommaso et al. 2017a) which orchestrates all processes and handles the retrieval and execution of the container, thereby ensuring seamless portability and reproducibility across platforms.

The pipeline generates a results directory for each sample, named accordingly and organized as follows:

- The *fastQC* directory contains the quality checks for each filtering stage.
- The *assembly_spades* directory contains the final assembly (*contigs.fasta*) not filtered as well as other internal files generated by *SPAdes*, useful mainly for debugging or further exploration of the assembly.
- *_contigs.filtered.fasta*: the final assembly file obtained after applying the length-based filtration step (according to the minimum length parameter).
- *_diamond_blastx.tsv:* Raw output file generated by *Diamond BLASTx*.
- *_contigs_read.tsv*: a tab delimited summary listing all the contigs
- *_all_viral_taxonomy.txt*: raw tab-delimited file associating each assembled contig with its corresponding viral taxonomic assignation.
- *_taxonomy_viral.clean.txt*: a cleaned and aggregated tab-delimited table reporting read counts per viral taxon to estimate relative abundance.
- *_contigs_reads.tsv*: alignment summary file reporting the number of reads mapped to each contig.
- *_rmrdna.fastq.gz*: filtered read file containing sequences retained after quality filtering, trimming (if enabled), and removal of ribosomal DNA contamination.

### Experimental infection and sample preparation

To validate the pipeline, we used four experimentally infected mosquito samples with known viral compositions. Females of *Aedes albopictus* (Strain la Lopé, Gabon) were experimentally infected with 1×10^7^ PFU/mL for each of the six viruses tested (Table 1) and incubated for 14 days post blood feeding following a previous protocol (Jiolle et al. 2021). After incubation, females were killed and stored at -80°C. Pools of five females per virus were prepared and then used for RNA extraction. Whole mosquitoe’s tissues were homogenized in 100µL of lysis buffer and 1.0mm glass beads (Biospec products). RNA fragments were purified from each sample using the Nucleospin RNA extraction kit (Macherey-Nagel) according to manufacturer recommendations. Purified RNA was quantified using Qubit HS RNA (Thermo Fisher Scientific).

**Table 1:**
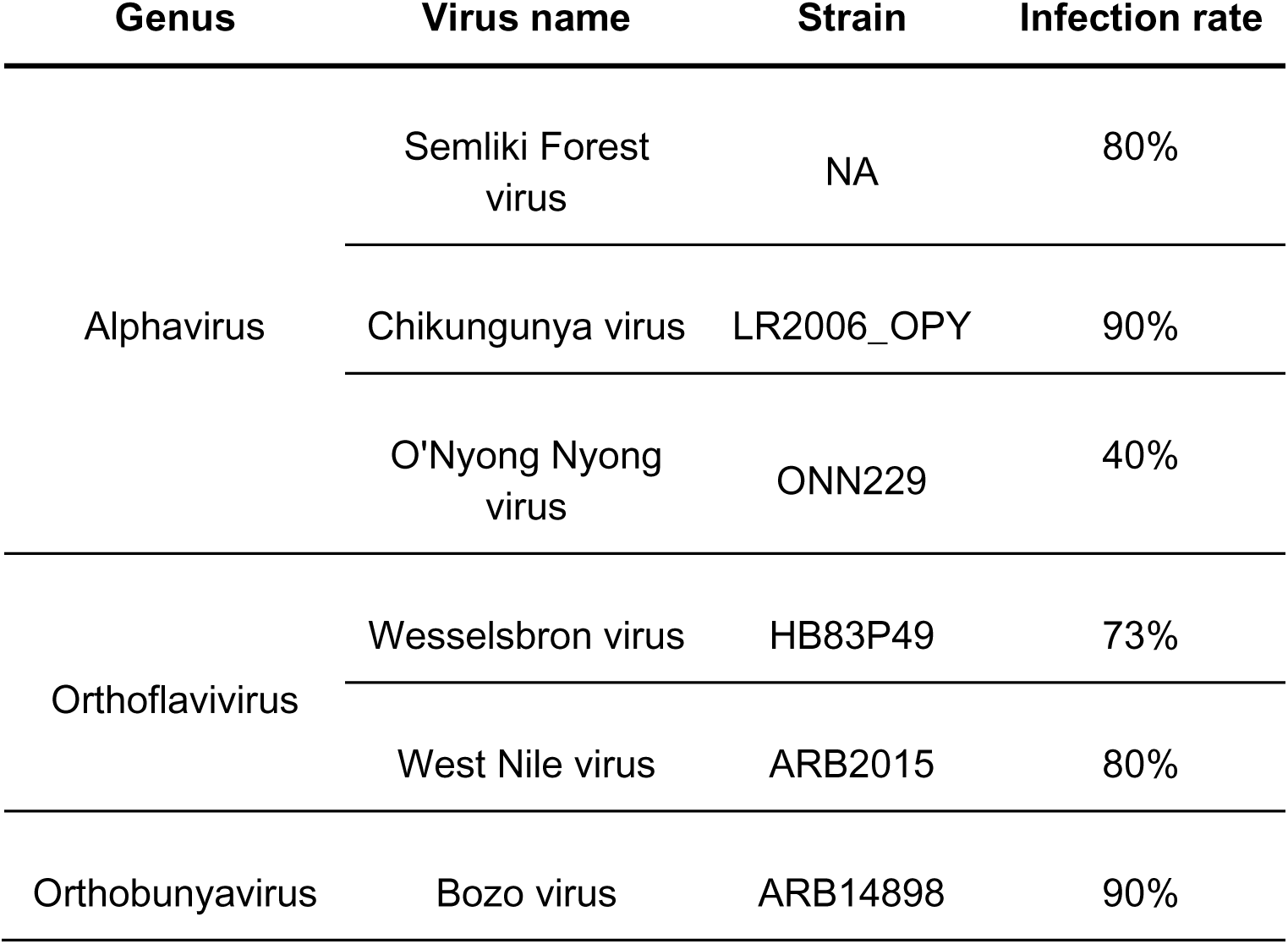
Viruses used for experimental infections, strains and infection rate. The infection rate corresponds to the percentage of positive females by RTqPCR. NA: not available.

To simulate realistic metagenomic samples and introduce biological background noise, a separate RNA extract was prepared from a pool of 30 uninfected *Aedes aegypti* females (Paea, strain, Tahiti, French Polynesia). *Aedes aegypti* was chosen as a source of background RNA because large laboratory-reared colonies were readily available, ensuring a homogeneous and infection-free material. This strategy aimed at reproducing the biological noise usually encountered in field-collected samples, where viral RNA typically represents only a minor fraction of total RNA.

Three successive dilutions of the RNA extraction from infected mosquitoes were prepared. The first mixture (Mix A) was obtained by combining 4 µL from each pool corresponding to the six viral infections (Semliki Forest virus, Chikungunya virus, O’nyong-nyong virus, Bozo virus, Wesselsbron virus and West Nile virus). A 1:100 dilution of Mix A generated Mix B, and a subsequent 1:100 dilution of Mix B yielded Mix C. Each of these dilutions was then supplemented with *Aedes aegypti* RNA in order to mimic realistic biological samples (see Supplementary Information 1). This resulted in three final sample types: MixA sample, consisting of 18 µL of Mix A combined with 6 µL of mosquito RNA; MixB sample, containing 18 µL of Mix B and 6 µL of mosquito RNA; and MixC sample, prepared with 18 µL of Mix C and 6 µL of mosquito RNA.

An additional sample (AltMix sample) consisting of a pool of eight *Ae. albopictus* specimens coinfected with Aedes albopictus densovirus 2 (strain AalDV2) and chikungunya virus were prepared. The presence of both viruses in this pool was confirmed by RT-qPCR (Supplementary information 2). This sample enabled us to assess the pipeline’s capacity to detect this viral mix, including Aedes albopictus densovirus 2 (strain AalDV2). While the other viruses tested are RNA viruses, AalDV2 is a single-stranded DNA virus classified within the family *Parvoviridae* and *Brevihamaparvovirus dipteran 2* species. The preparation of samples and mock viromes using experimental infections is illustrated in Figure 2.

**Figure 2:**
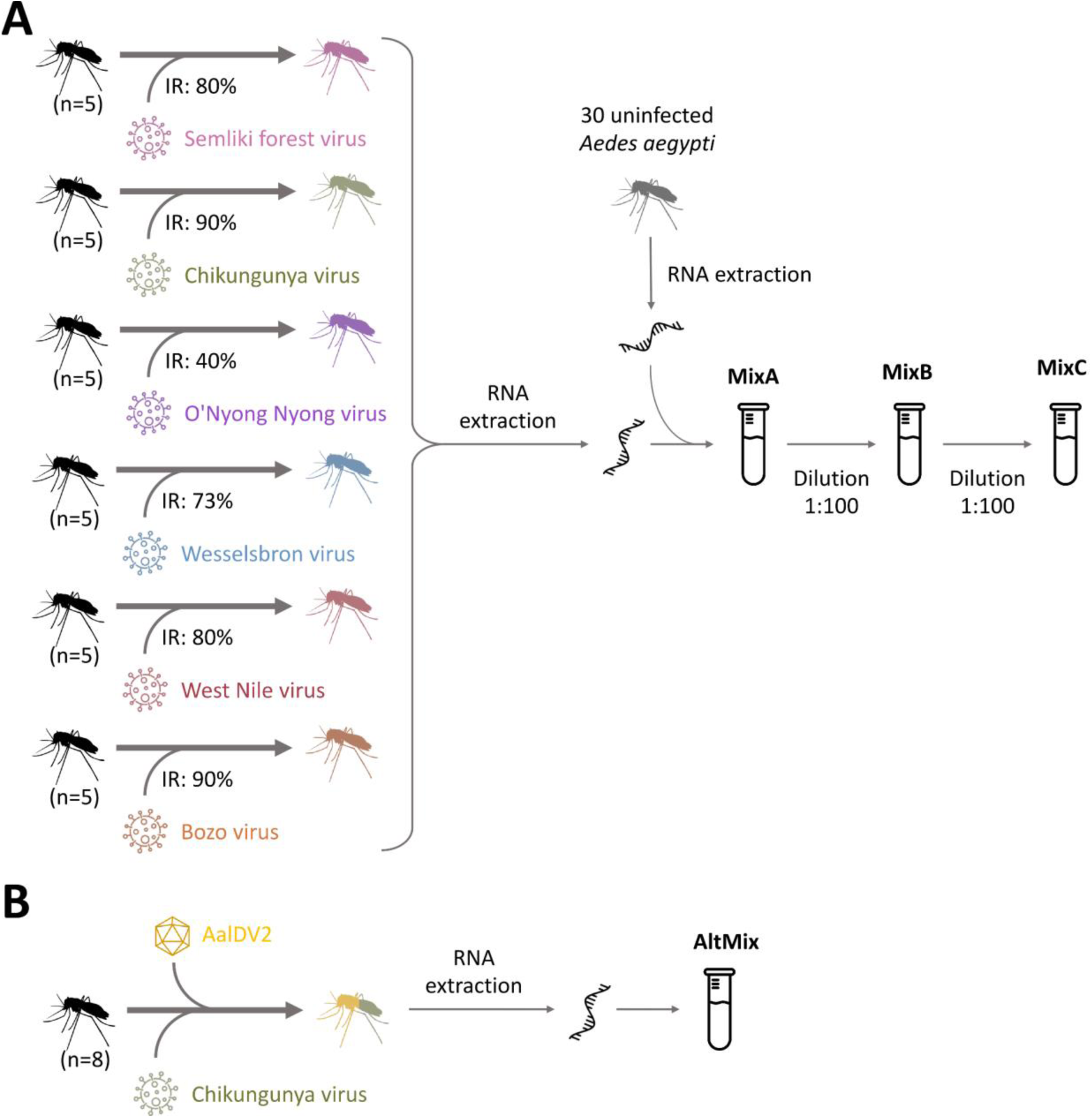
Simplified preparation of the experimental samples. **A.** Preparation of samples MixA, MixB and MixC. The mosquitoes (shown in black) are experimentally infected female *Aedes albopictus*. For each experimental infection, the infection rate (IR) is specified. **B.** Preparation of sample AltMix. The mosquitoes (in black) are experimentally infected female *Aedes albopictus*. AalDV2: Aedes albopictus densovirus 2.

### Libraries preparation and target enrichment

The RNA was converted to a first-strand cDNA using Superscript III and the second strand was synthesized using NEBNext® Ultra™ II Non-Directional RNA Second Strand Synthesis Module (E6111), followed by a purification using AMPure Xp beads (Beckman Coulter) and Qubit quantification (HS RNA).

For MixA to MixC samples, library preparation was performed following the SureSelect xt Target Enrichment System protocol (Supplementary information 1). Due to issues during library preparation, the following libraries and AltMix samples were performed using Twist protocol (Supplementary information 2). Twist viral probe panel (Twist Bioscience) was designed using CATCH (Metsky et al. 2019) to target 533 virus sequences from 15 families, as described in Supplementary Table 2. After verifying that the amount of DNA was sufficient for all pools, the pools were clustered to generate 240 million raw reads on Novaseq6000 with 2 × 150 base pairs (bp) paired-end reads (IntegraGen).

### Generation of the simulated data

Simulated paired-end metagenomic reads have been generated using *CAMISIM v1.3* (Quick et al. 2016) as in Mastriani et al. (2022). This simulation was designed to provide a controlled and reproducible framework to evaluate specific aspects of the pipeline, including the detection of viral sequences at different abundance levels and the ability to filter out non-viral contaminant sequences (Antipov et al. 2020). Three viral sequences were selected: Hepatitis A virus (NC_001489.1) and Ippy virus segment S (NC_007905.1) and L (NC_007906.1). The bacterial genome of *Helicobacter hepaticus* (AE017125.1) was used as a contaminant. The inclusion of a bacterial genome as a dominant component was intended to simulate a challenging background and to assess the robustness of contamination filtering and taxonomic assignment steps. This design does not aim to fully replicate a specific biological system, but rather to test the pipeline under stringent and controlled conditions. We simulated 10 metagenomic samples as paired-end sequencing from Illumina HiSeq 2500 (the ART Illumina model) with a mean fragment length of 270 bp. The abundance distribution was created by sampling from a log-normal distribution with default parameters. Among the ten generated samples, we selected the one that provided the most challenging and informative test conditions (Simulation 9 in Supplementary Table 3). In this sample, bacterial contamination (intended to be filtered out by the pipeline) was predominant. Two viral sequences were present in balanced proportions (Hepatitis A virus and Ippy virus segment S), while a third was included at a lower abundance (Ippy virus segment L).

### ViroSeek pipeline execution

Validation of ViroSeek v0.0.2 was performed using one simulated metagenome (SIM sample) and four samples from mosquitoes experimentally infected with known viruses (MixA, MixB, MixC and AltMix samples). The *Nextflow* pipeline (with *Nextflow v24.10.4*) was executed on all five control samples. Reads pre-processing was performed using *trimgalore*. To minimize erroneous taxonomic assignments associated with short contigs, a minimum contig length of 140 bp was applied to the assembly. Additionally, stringent alignment parameters were used, including a maximum *e-value* of 1e-5, a minimum percent identity of 96.0%, and a minimum query coverage of 51%. Following the *Diamond* documentation, the *--fast* option was used since the minimum percent identity exceeded 90%. These parameters were defined in order to identify all target species, while remaining as stringent as possible to avoid false positives. The database used for taxonomic classification was derived from the National Center for Biotechnology Information Reference Sequence non-redundant protein database (Pruitt et al., 2005) and was not restricted to viral sequences, unlike the default configuration implemented in ViroSeek. We selected a database not restricted to viral sequences in order to facilitate the detection of potential false positives, particularly when comparing results with other pipelines. It should also be noted that lower e-value thresholds are recommended when using smaller databases to reduce the occurrence of false-positive assignments.

### Comparison between existing pipelines and ViroSeek

To assess the performance and suitability of ViroSeek, we compared it with several existing pipelines commonly used for viral metagenomic analysis: Taxprofiler v1.2.4 (Stamouli et al. 2023), MetaDenovo v1.3.0 (Lundin et al. 2025), Viralmetagenome v1.1.1 (Klaps et al. 2026), and VirusTaxo v2 (Github last accessed in October 2025; Raju et al. 2022). Comparison with MicroSeek could not be performed because the pipeline’s code and workflow are not openly accessible. Furthermore, for PIMGAVir, we were unable to run the pipeline due to excessive RAM requirements.

For each test sample generated in this study, we compared the performance of different pipelines by evaluating computational resource usage (CPU time and maximum memory) and sensitivity toward various virus species. Taxprofiler,MetaDenovo and Viralmetagenome, were executed directly on the raw sequencing reads, whereas VirusTaxo was applied to the assemblies generated by ViroSeek, as described above. For Taxprofiler and MetaDenovo, the same NCBI database as ViroSeek was used, and Diamond served as the taxonomic classifier. Taxprofiler, MetaDenovo and Viralmetagenome are from the nf-core collection of workflows (Ewels, Magnusson, Lundin, and Käller 2016), utilising reproducible software environments from the Bioconda (Grüning et al. 2018) and Biocontainers (da Veiga Leprevost et al. 2017) projects. They were executed with *Nextflow v25.10.0* (Di Tommaso et al. 2017b).

For Taxprofiler, data was processed using *nf-core/taxprofiler v1.2.4* (Stamouli et al. 2023), and was configured to perform quality control, including read merging, minimum length filtering at 30 bp, and duplicate removal. Short-read complexity assessment was also enabled. Sequencing quality control and short read preprocessing were respectively performed with *FastQC* (Andrews et al. 2023) and fastp (Chen et al. 2018; Chen 2025)c classification was carried out with *Diamond* (Buchfink et al. 2021). Pipeline results statistics were summarised with *MultiQC* (Ewels, Magnusson, Lundin, and Käller, Max 2016).

For MetaDenovo, data was processed using *nf-core/metadenovo v1.3.0* (doi: 10.5281/zenodo.10666590) More precisely in this workflow, raw reads were quality-checked using *FastQC* (Andrews et al. 2023) and trimmed with *TrimGalore* (Krueger et al. 2023). Reads were aligned using *BBMap* (https://sourceforge.net/projects/bbmap/), and alignment files were processed with *Samtools* (Danecek et al. 2021). Sequences were filtered to remove potential ribosomal DNA contamination using the same SILVA rRNA database as with ViroSeek (Quast et al. 2013, release 138). De novo assembly was performed using *SPAdes* (Prjibelski et al. 2020). Gene prediction was conducted using Prodigal (Hyatt et al. 2010), and taxonomic annotation was performed with *Diamond* (Buchfink et al. 2021), with lineage assignment using *Taxonkit* (Shen and Ren 2021). Gene abundance was quantified using *featureCounts* from *Subread* (Yang et al. 2014). Pipeline results statistics were summarised with *MultiQC* (Ewels et al. 2016). For both pipelines, we kept only results with a minimum *e-value* of 1e-5.

For Viralmetagenome, data was processed using *nf-core/viralmetagenome v1.1.1* (Klaps et al. 2026). Assembly of reads was conducted using *SPAdes* (Prjibelski et al. 2020), with a minimum contig size threshold of 140 bp. Sequence clustering was performed using the *MMseqs2* clustering method (Kallenborn et al. 2025). Variant calling and annotation steps were omitted from the workflow.

For each pipeline, the parameters were selected to closely align with those used in ViroSeek, thereby ensuring that any observed differences could be attributed primarily to pipeline performance rather than to discrepancies in software configuration.

### Post-pipelines downstream analysis

After completion of the pipelines, a post-processing workflow was applied to harmonize taxonomic assignments, generate abundance tables and downstream analyses. All viral taxonomy output files generated (\**_taxonomy_viral.clean.txt*) were first collected from individual sample directories and renamed to a standardized *krona* format. These files were concatenated into a single taxonomy file, allowing a unified representation of viral taxa across samples. Taxonomic tables were then reconstructed in *R v4.2.3* (R core Team 2023) and formatted to comply with the International Committee on Taxonomy of Viruses (ICTV) hierarchical structure. Only entries belonging to the Viruses realm were retained for downstream analyses. Read count data were aggregated per taxonomic assignment using a custom Python script modified from Hayer *et al*. (2023), producing an abundance table summarizing the total number of reads per taxon and per sample. This table was combined with the curated taxonomy and sample metadata to construct a *phyloseq* object, from package *phyloseq v1.42.0* (McMurdie and Holmes 2013), in *R v4.2.3* (R core Team 2023). Relative abundances were calculated by normalizing read counts to percentages within each sample. Taxa were agglomerated at the species level, and bar plots were generated to visualize viral community composition with *ggplot2* v3.5.1 (Wickham 2016). Target viral genera contained in the probes panel were specifically extracted for graphical representation for clarity.

## Results

### Pipeline validation

The viral panel used in this study includes representative arboviruses of medical relevance as well as a DNA virus belonging to a distinct viral family, allowing evaluation across different genome types commonly encountered in surveillance datasets. Although this benchmarking provides a controlled and realistic evaluation of pipeline performance, it does not encompass the full diversity of known viral taxa, particularly highly divergent or under-represented viruses. Across five control datasets, ViroSeek detected 100% of expected viral species, including low-abundance targets, while maintaining low false positive rates. It outperformed Taxprofiler, MetaDenovo and VirusTaxo in both computational efficiency (approximately 4 and 20 times faster respectively) and sensitivity for virus detection. To validate the ViroSeek pipeline, we designed five samples containing known viral targets, belonging to *Alphavirus*, *Orthoflavivirus*, and to DNA viruses. These samples covered different levels of complexity, from dilution series of viral mixes (MixA–MixC) to alternative compositions combining RNA and DNA viruses (AltMix, SIM). This setup provided controlled conditions to assess detection sensitivity and specificity.

The analysis found all the viruses expected in MixA (6/6) and MixB (6/6) samples, confirming ViroSeek’s ability to detect a wide range of viruses in complex samples (Figure 3). In the most diluted condition (MixC) all expected viruses were still detected (6/6), however less species were detected overall and a ∼20% reduction in relative abundance of enriched families was observed compared to MixA and MixB, indicating a sensitivity limit of the pipeline under low-abundance conditions. The target viruses were also correctly detected in SIM sample (2/2). In AltMix sample, chikungunya virus was the dominant virus detected and the presence of Anopheles gambiae densovirus (AgDV) was also reported (2/2). However, its low abundance prevented direct visualization in the graphs, but it is symbolized by arrows (Figure 3). The underrepresentation of AgDV might be attributed to the biology of densoviruses, whose single-stranded DNA genome is only captured during the replication phase, via the expression of viral transcripts (Kimmick et al. 1998). It is important to note that AgDV belongs to the family *Parvoviridae*, species *Brevihamaparvovirus dipteran 1*, and not to *Brevihamaparvovirus dipteran 2* as reported by ViroSeek. This taxonomic misassignment will be further discussed below. Furthermore, Orthoflaviviruses not introduced during experimental infection were detected in MixA sample as Sepik virus, Rift Valley virus, cell fusing agent virus and Aedes flavivirus. The identification of Sepik virus and Rift Valley virus corresponds to a taxonomic misassignment, while Cell fusing agent virus and Aedes flavivirus are insect-specific flavivirus commonly found in mosquito samples (Bolling et al. 2012; Fu et al. 2025). With the exception of the cell fusing agent, which remains detectable in all samples. Other viruses were present at low abundance and are no longer detectable after successive dilution steps. In addition, the pipeline cleaning steps effectively removed contaminating sequences, such as bacterial sequences in SIM sample (Table 2) and host sequences in the other samples (Figure 3).

**Figure 3:**
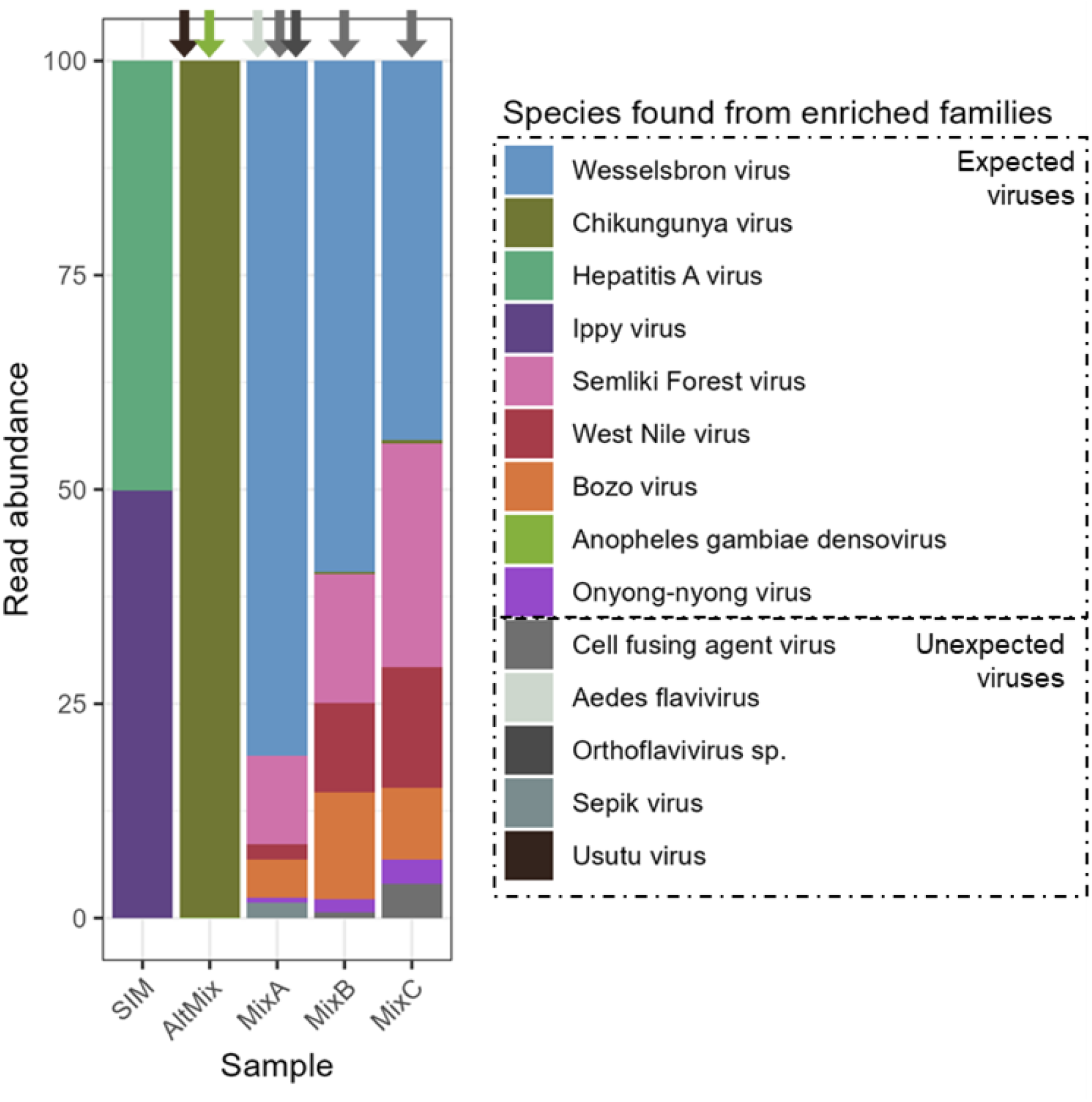
Viral relative abundance from targeted families. Only viruses belonging to the targeted families are shown, all other viral taxa from non-targeted families are excluded for clarity. “Unexpected viruses” refer to additional viruses detected that were not the species used in the experimental infections but belonged to viral families targeted by the enrichment probe panel (Supplementary Table 2). Abundance corresponds to the percentage of sequencing reads mapped to each viral species. Coloured arrows correspond to viral species detected in the samples but not visible on the graph due to low abundance.

**Table 2:** Summary of viral genomes reconstruction for SIM sample. . Included reference genome, expected genome size, contig length and corresponding read depth.

| Reference genome | Reference length (bp) | Taxonomic assignation | Contig length (bp) | Number of reads mapped |
| --- | --- | --- | --- | --- |
| Ippy virus segment S (NC_007905.1) | 3366 | Ippy virus segment S (YP_516231.1) | 3314 | 370 |
| Ippy virus segment L (NC_007906.1) | 7316 | Ippy virus segment L (YP_516233.1) | 278 | 3 |
| Hepatitis A virus (NC_001489.1) | 7478 | Hepatitis A virus (NP_740559.1) | 586 | 25 |
|  |  | Hepatitis A virus (NP_041007.1) | 6798 | 350 |
| *Helicobacter hepaticus* (AE017125.1) | 1799146 | - | - | - |

Two cases of taxonomic misassignment were observed. In AltMix sample, two contigs were assigned to AgDV, while we expected to detect AalDV2. Inspection of the sequences reveals that the strain has been correctly assigned to the capsid protein Q90187.2 (from AalDV2), but this protein has been linked to the wrong species (AgDV). This misassignment is likely due to an inconsistency in the NCBI database, where this sequence is referenced as AgDV rather than AalDV2.

In MixA sample, two contigs were falsely assigned to Sepik virus, a virus not included in the experimental mixture. After verification, the assignment was explained by a protein homology with Wesselsbron virus, preventing reliable discrimination between the two. It was not possible to increase the stringency of the filters without affecting the detection of other target viruses. Finally, a 1948bp contig assigned to the Usutu virus was detected in the AltMix sample, although no Usutu virus had been used in the preparation of this library. However, this sample had been prepared in parallel and pooled for sequencing with other Usutu-positive libraries as part of another project on *Culex* mosquitoes. As only a single contig was assigned to this virus, and considering the library preparation context, this isolated detection suggests potential cross-contamination during preparation or pooling of libraries. This observation does not call into question the efficiency of the pipeline, but rather highlights the importance of rigor in the upstream stages of the experimental protocol, particularly when handling enriched libraries. It also emphasizes the need to interpret certain signals with caution and, when biological confirmation is required, to validate them using complementary approaches.

### Comparison of ViroSeek efficacy and sensitivity with existing pipelines

To evaluate the efficacy and sensitivity of ViroSeek, we compared its performance with that of existing metagenomic pipelines, focusing on its ability to detect viruses across diverse test samples.

Firstly, in terms of computational performance, ViroSeek required approximately 184 CPU hours to process the five test samples, compared to 3,891 for Taxprofiler, 453 for MetaDenovo, and 295 for Viralmetagenome. Regarding memory usage, ViroSeek utilized a maximum of 42.3 GB RAM, whereas Taxprofiler, MetaDenovo, and Viralmetagenome required 40.3, 42, and 72 GB respectively (Figure 4). Thus, ViroSeek was the fastest pipeline, and its memory consumption was equivalent to Taxprofiler and MetaDenovo.

**Figure 4:**
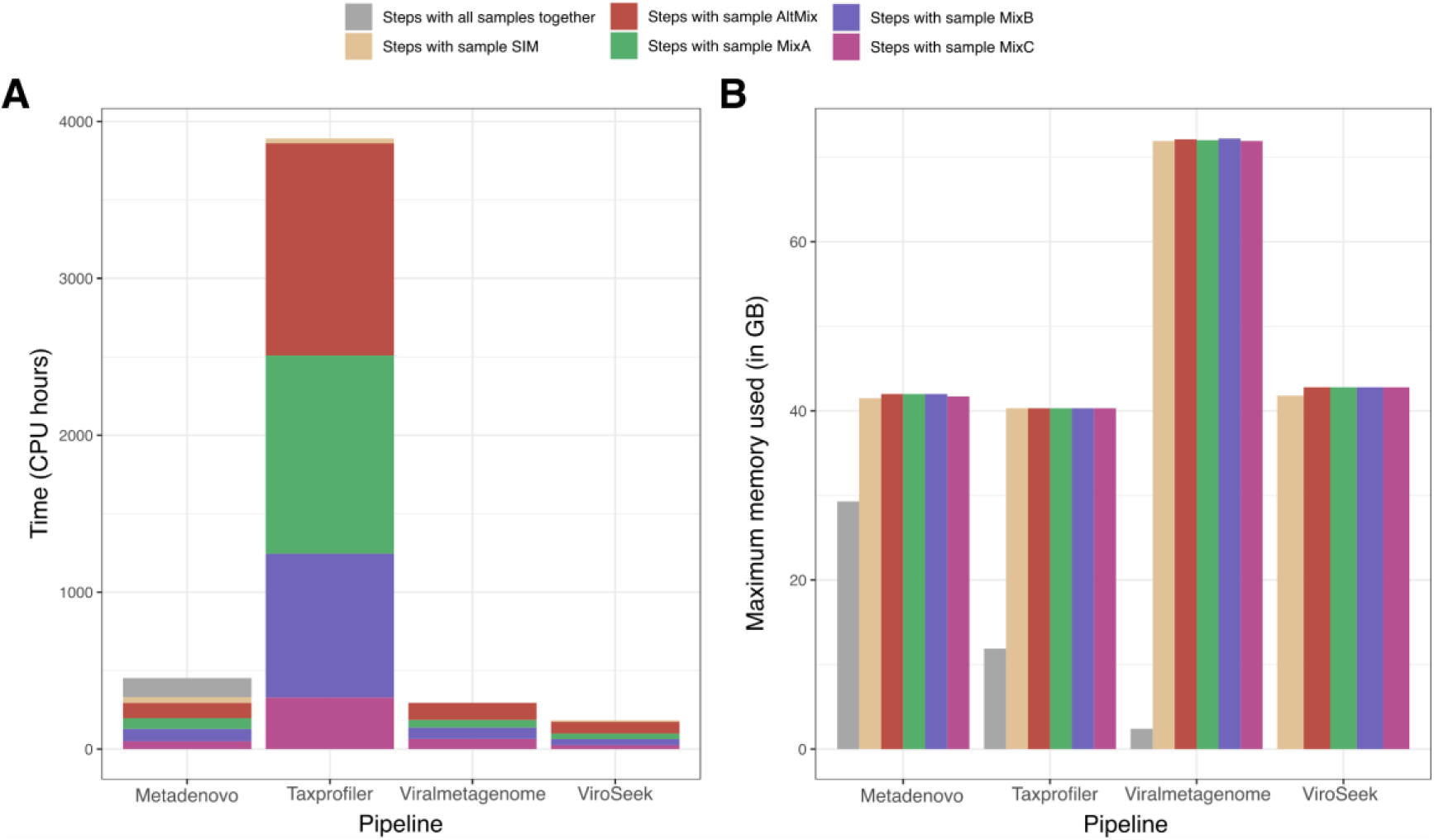
Comparison of the performance of the different pipelines tested. **A.** Bar chart showing the computational time required by each pipeline, expressed in CPU hours. **B.** Bar chart presenting the maximum memory usage (in GB) for each pipeline. In both plots, colors correspond to the individual samples processed. For MetaDenovo, Taxprofiler and Viralmetagenome, some steps operate on all samples together: it is represented in gray.

ViroSeek successfully detected all expected viruses in every dataset, including the simulated sample (SIM), AltMix, and the three mixed samples (with all dilutions from 1 to 1:10000). Similarly, TaxProfiler and ViralMetagenome recovered all expected viruses in all datasets (Table 3). Both pipelines correctly identified the expected densovirus species (*Aedes albopictus* densovirus 2). However, TaxProfiler additionally assigned a subset of reads to *Anopheles gambiae* densovirus, reflecting ambiguity in species-level assignment. In contrast, MetaDeNovo showed reduced sensitivity, and detected all expected viruses only in the simulated sample. In AltMix, only Chikungunya virus was detected among the expected viruses, whereas four of the six expected viruses were recovered in MixA and MixB, with Semliki Forest virus and Chikungunya virus consistently missed. In MixC, only three expected viruses were detected, with Semliki Forest virus, Chikungunya virus, and Wesselsbron virus not recovered. VirusTaxo exhibited the lowest sensitivity, detecting only one expected virus in the simulated sample, none in AltMix, and three of the six expected viruses in MixA, B and C.

**Table 3.** Summary of the sensitivity comparison between ViroSeek and the other pipelines evaluated. For each sample, the expected number of viral species was defined according to the experimental and bioinformatic protocols (see Materials and Methods). For each pipeline, we report both the number of expected viral species correctly detected and the total number of unexpected viral species identified. “Unexpected viruses” refer to additional viruses detected that were not the species used in the experimental infections but belonged to viral families targeted by the enrichment probe panel (Supplementary Table 2). “Confirmed false positive” are viruses species that are “unexpected viruses” but they are not reported to be naturally part mosquito virome, so they are true confirmed false detections.

|  |  | SIM | AltMix | MixA | MixB | MixC |
| --- | --- | --- | --- | --- | --- | --- |
|  | Number of expected viruses | 2 | 2 | 6 | 6 | 6 |
| **ViroSeek** | Number of expected viruses found | 2 | 2 | 6 | 6 | 6 |
|  | Number of unexpected viruses | 0 | 7 | 16 | 20 | 16 |
|  | Confirmed false positive | 0 | 0 | 2 | 0 | 1 |
| **Metadenovo** | Number of expected viruses found | 2 | 1 | 4 | 4 | 3 |
|  | Number of unexpected viruses | 0 | 28 | 14 | 14 | 37 |
|  | Confirmed false positive | 0 | 13 | 1 | 0 | 2 |
| **Taxprofiler** | Number of expected viruses found | 2 | 2 | 6 | 6 | 6 |
|  | Number of unexpected viruses | 1 | 40 | 160 | 144 | 36 |
|  | Confirmed false positive | 1 | 18 | 79 | 68 | 8 |
| **VirusTaxo** | Number of expected viruses found | 1 | 0 | 3 | 3 | 3 |
|  | Number of unexpected viruses | 0 | 2 | 3 | 2 | 2 |
|  | Confirmed false positive | 0 | 0 | 0 | 0 | 0 |
| **Viralmetagenome** | Number of expected viruses found | 2 | 2 | 6 | 6 | 6 |
|  | Number of unexpected viruses | 2 | 21 | 65 | 65 | 42 |
|  | Confirmed false positive | 2 | 12 | 23 | 26 | 29 |

The number of unexpected viral detections differed substantially among pipelines (Table 3). ViroSeek produced no unexpected detection in the simulated dataset. The number of unexpected species increased in the three mixed samples, reaching 7, 16, 20 and 16 detections in AltMix, MixA, MixB and MixC, respectively. Aside from the Sepik virus and Rift Valley virus assignment, which has been confirmed to be a false positive, most unexpected viruses detected by ViroSeek corresponded to associated viruses of mosquitoes or viruses plausibly linked to the mosquito microbiome. These included insect-specific viruses such as Aedes binegev-like virus 1, Culex picorna-like virus 1, Aedes flavivirus and cell-fusing agent virus (Hoshino et al. 2009; Sadeghi et al. 2018; Cholleti et al. 2018; Guarido et al. 2021; Parry et al. 2021). Additional detections included bacteriophages associated with bacterial taxa commonly found in the mosquito microbiome, including members of the *Herelleviridae*, *Ackermanviridae*, *Microviridae* and *Siphonoviridae* families, as well as phages infecting *Klebsiella* spp. or other bacterial groups known to colonize mosquitoes (Barylski et al. 2020; Kang et al. 2020; Mosquera et al. 2023; Nadeem et al. 2026). Metadenovo generated substantially more unexpected viral detections, with 2, 28, 14, 14, 37 unexpected species identified in the SIM, AltMix, MixA, MixB, MixC samples, respectively. Several of these assignments corresponded to arboviruses that were not present in the samples, including getah virus, Sindbis virus, Ross River virus, Mayaro virus, Venezuelan equine encephalitis virus and Middleburg virus. Similar patterns were observed with Taxprofiler, which detected 1 unexpected species in the simulated dataset, 40 in AltMix, 60 in MixA, 144 in MixB and 36 in MixC. In the simulated dataset, the *Mammarenavirus* sequence was misclassified as Kitale virus, a related member of the same viral genus. In the experimental datasets, numerous unexpected arboviruses were detected, including Mayaro virus, getah virus, Ross River virus, yellow fever virus or Zika virus.

Although VirusTaxo produced relatively few unexpected species assignments (0 in SIM, 2 in MixB and MixC and 3 in Mix A and C), these results should be interpreted cautiously because the majority of viral reads remained unclassified at species level. More than 90% of viral reads remained unclassified at species resolution. In the simulated dataset, hepatitis A virus accounted for approximately 50% of all reads but could not be assigned at the species level (Figure 5). Several contigs were correctly assigned at the family or genus level but lacked sufficient resolution for species identification, consistent with previous reports (Raju et al. 2022). ViralMetagenome detected all expected viruses but generated more unexpected detections than ViroSeek, with 2, 21, 65, 65, 42 unexpected species in the SIM, AltMix, MixA, MixB and MixC, respectively. Unexpected assignments included several arboviruses, such as West Niles virus, o’nyong-nyong virus, Semliki Forest virus in AltMix and dengue virus, Saint Louis encephalitis virus, yellow virus or Ross River virus for MixA, B and C.

**Figure 5:**
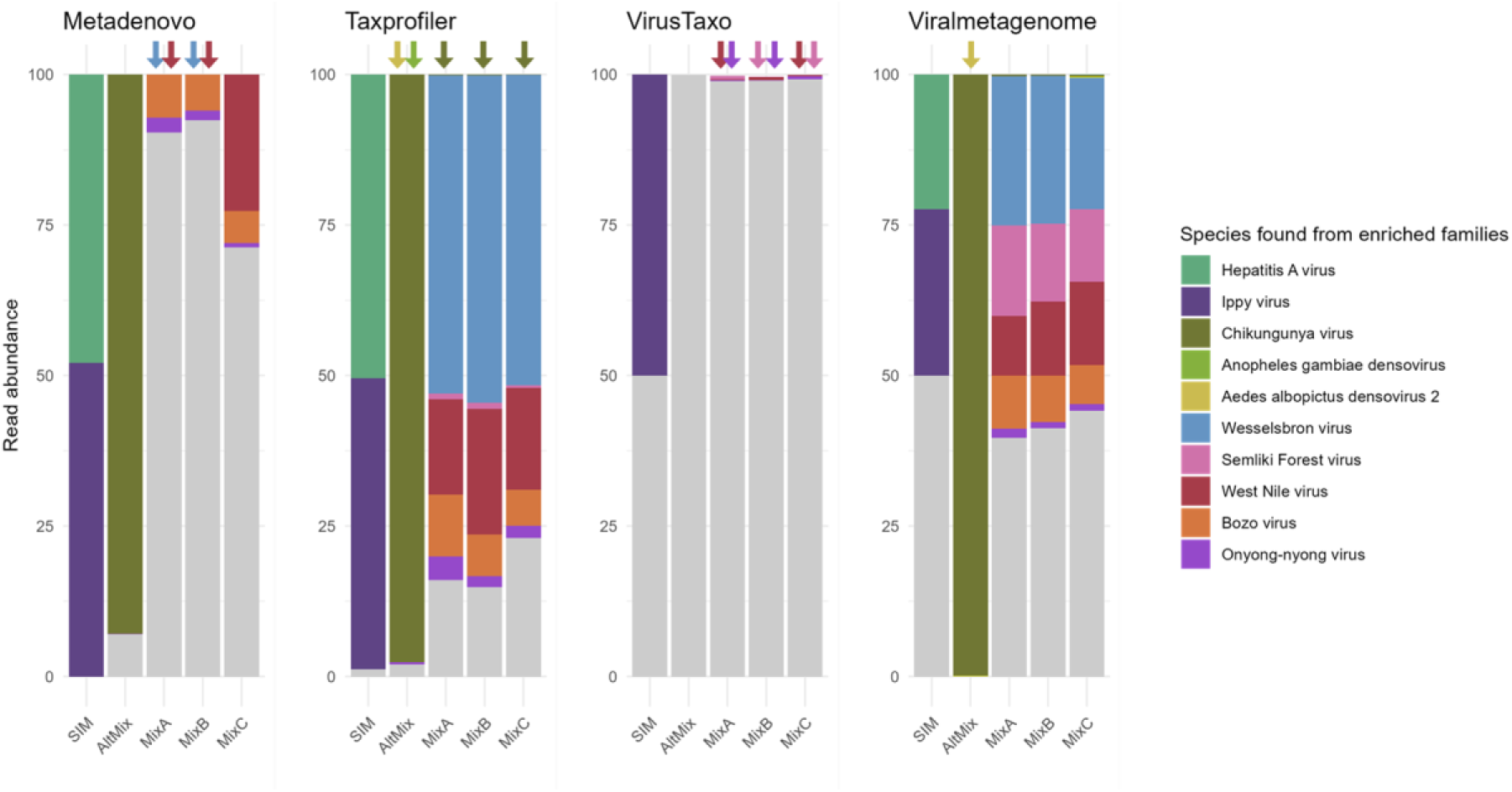
Viral relative abundance from targeted families across alternative pipelines. Only viruses belonging to the targeted families are shown, all other viral taxa are excluded for clarity. “Other” refers to species not used in the experimental infections but that belonged to viral families targeted by the enrichment probe panel (Supplementary Table 2). Due to the high number of false positives, these signals were grouped into a single category to simplify the interpretation of the figure. Coloured arrows correspond to viral species detected in the samples but not visible on the graph due to low abundance.

Overall, while all pipelines, except Metadenovo and VirusTaxo, achieved high sensitivity for the detection of expected viruses, important differences were observed in the number and nature of unexpected viral assignments (Figure 5). Unexpected detections generated by ViroSeek were predominantly composed of insect-specific viruses and bacteriophages plausibly associated with the mosquito microbiome, whereas competing pipelines frequently assigned reads to medically relevant arboviruses that were not present in the samples. These results indicate that ViroSeek maintains high sensitivity while reducing biologically implausible taxonomic assignment relative to the other evaluated pipelines.

## Discussion

Several virome analysis pipelines already exist, as detailed in the introduction, but many of them are either highly specialized, complex to install or use, or poorly maintained. ViroSeek addresses this gap by providing a user-friendly workflow optimized for virus-enriched libraries, making it accessible to non-expert users while maintaining robust performance. Beyond producing reliable taxonomic assignments, the pipeline also generates assembled contigs that can be reused for phylogenetic analyses, supporting comprehensive evolutionary and comparative studies (Kane et al. 2024). During validation, ViroSeek consistently identified all expected viruses, even with high viral complexity or low-abundance targets, including single-stranded DNA densoviruses, which are only efficiently detected during the replicative phase when viral transcripts are produced. ViroSeek was not only faster but also provided more accurate taxonomic assignments than the other tools evaluated in this study (Figures 4 and 5). These characteristics make it a particularly relevant and effective tool for the analysis of viral metagenomes, especially in the context of arbovirus research.

In the present study, the interpretation of unexpected viral detections requires caution because the complete virome of whole mosquitoes is not fully characterized. Consequently, unexpected viral species cannot systematically be considered as false positives. Many of the other taxa detected by ViroSeek corresponded to insect-specific viruses previously reported in mosquitoes or to bacteriophages potentially associated with members of the mosquito microbiome. The biological relevance of these detections therefore cannot be excluded. However, we detect only two false positives (Rift Valley virus and Sepik virus) in all samples and one assignment of contigs to incorrect taxa (e.g AgDV instead of AalDV2) in AltMix highlights the limitation of current taxonomic databases. Such misassignments are not necessarily attributable to ViroSeek itself, but rather to the lack of sufficiently discriminant markers; when viral sequences are highly conserved between closely related taxa, taxonomic classification becomes uncertain. This shows the importance of manual validation for critical cases, as well as the need for carefully curated databases and the development of more specific probes to improve discrimination between related viral species. In contrast, several competing pipelines frequently assigned reads to medically important arboviruses that were absent from the experimental design, including yellow fever virus or Zika virus. Such assignments cannot possibly originate from the virome of the mosquitoes and are therefore considered taxonomic classification errors. These results suggest that ViroSeek not only achieves high sensitivity for the detection of expected viruses but also reduces false assignment compared with other evaluated pipelines. This characteristic is particularly important for arbovirus surveillance, where false identification of medically relevant viruses might mislead epidemiological interpretations.

Another critical point highlighted concerns the quality of upstream experimental steps. The detection of a contig assigned to the Usutu virus in a sample that did not experimentally contain it, highlights the risk of cross contamination during library pooling, especially with high viral concentrations. Even if this result does not invalidate the performance of the pipeline, it does illustrate the importance of rigorous control of laboratory practices. ViroSeek’s effectiveness has been empirically validated under predefined conditions, in particular with regard to taxonomic assignment parameters such as range culling, minimum query cover and identity thresholds. These parameters have been optimized to guarantee a compromise between sensitivity and accuracy. Reducing these thresholds to increase detection could expose users to an increased risk of false positives or erroneous assignments, particularly in the case of viruses that are genetically close or poorly referenced in databases. We therefore evaluated the impact of the sequence identity used for taxonomic assignment. Lowering the identity threshold from 96% to 90% increased the number of unexpected viral species detected across samples (Supplementary Figure 1). However, this lower threshold did not result in an increase in false positives. Instead, most additional detections corresponded to insect-associated viruses. Because the complete virome of mosquitoes remains incompletely characterized, it is difficult to determine whether these additional assignments reflect real viral diversity that would be missed under more stringent criteria or an increased rate of false positives. Consequently, no definitive conclusion can be drawn regarding the optimal identity threshold for all applications. To accommodate different research objectives, the identity threshold remains configurable by user. However, we recommend testing ViroSeek on control samples with known viral composition to calibrate thresholds and ensure result reliability. In the absence of such controls, the default parameters experimentally validated in this study should be used. Furthermore, as demonstrated by our results with Orthoflaviviruses, sample dilution is advisable when the objective is to characterize the overall virome composition, but should be avoided when targeting low-abundance viral families.

ViroSeek has only been evaluated on eukaryotic viruses, although the workflow is compatible with both DNA and RNA viruses. Consequently, its application to bacteriophage or broader bacterial virome studies may be limited, and specialised tools such as VIBRANT (Kieft et al. 2020), VirSorter2 (Guo et al. 2021) or VirFinder (Ren et al. 2017) may be more appropriate in these contexts. In addition, the pipeline was specifically developed for enriched sequencing libraries, and its performance on shotgun metagenomic datasets has not been extensively assessed. Its sensitivity in untargeted metagenomic applications may be lower than workflows specifically designed for shotgun metagenomics, such as EasyMetagenome (Bai et al. 2025). Moreover, to maintain an accessible workflow for users with limited bioinformatics expertise, the pipeline implements a single assembler (*rnaviral SPAdes*) and a single taxonomic assignment approach (*BLASTx*). Although alternative assemblers, including other *SPAdes* modes and *PenguiN*, were considered, the *rnaviral SPAdes* was selected because it provides a balanced compromise between assembly quality and robustness for viral datasets. Jochheim *et al*. (2024) have shown that the *rnaviral* option can recover more complete viral genomes with fewer mismatches compared with other *SPAdes* assembly modes. While *PenguiN* may generate longer contigs, *rnaviral SPAdes* produces more conservative and precise reconstructions (Jochheim et al. 2024). For taxonomic assignment, ViroSeek relies on *BLASTx*, which was selected because it combines the sensitivity of protein-level searches while avoiding the biases introduced by explicit ORF prediction (Buchfink et al. 2021; Pearson 2014).

One current limitation of ViroSeek concerns the handling of PCR duplicates. By default, duplicate removal is performed on a coordinate-based tool *samtools markdup*. In the context of viral metagenomic enriched libraries, independent reads sharing identical mapping positions may be incorrectly classified as PCR duplicates and removed. The use of unique molecular identifiers (UMI) during library preparation can help overcome this issue by enabling the identification of true amplification-derived duplicates in the downstream analysis. Incorporating support for UMI-based deduplication, for example with UMI-tools (Smith et al. 2017), represents a potential direction for future development of Viroseek. In the meantime, users may avoid biases associated with *samtools markdup* by running the pipeline with the *--skip_dedup* option.

In conclusion, ViroSeek provides a user-friendly, and reproducible workflow for the analysis of target-enriched metagenomic sequencing data. Across all datasets, it accurately detected the expected viruses, including low-abundance and phylogenetically diverse strains, with less erroneous taxonomic assignments compared with other commonly used pipelines. In addition to taxonomic identification, the generation of assembled contigs enables downstream phylogenetic and comparative analyses. By facilitating standardized and reproducible virome analyses, ViroSeek has the potential to support arbovirus surveillance and studies of viral diversity across a wide range of ecological and epidemiological contexts.

## Acknowledgment

The authors acknowledge the ISO 9001-certified IRD i-Trop HPC (member of the South Green Platform) at IRD Montpellier for providing HPC resources that contributed to the research results reported in this paper. URL: https://bioinfo.ird.fr/- http://www.southgreen.fr. The bioinformatics analyses were also performed on the Core Cluster of the Institut Français de Bioinformatique (IFB) (ANR-11-INBS-0013). We would also like to thank James Fellows Yates for his assistance in addressing the resource limitations encountered with Taxprofiler. We would like to thank Mathilde Jaquet for helping us with the *phyloseq* package. This study received financial support by the Agence Nationale de la Recherche (ANR PRC TIGERBRIDGE, grant n° 16-CE35-0010-01 to CP). AB has been granted an IRD PhD Fellowship.

A preprint version of this article has been peer-reviewed and recommended by PCI Genomics (https://doi.org/10.24072/pci.genomics.100695).

## Data and code availability

The data for this study have been deposited in the European Nucleotide Archive (ENA) at EMBL-EBI under accession number PRJEB93907.

All the scripts, the related metadata and documentation are available in the DataSuds repository (IRD, France) at https://doi.org/10.23708/LTXIKA. Related documentations are openly available and granted under CC-BY license.

The pipeline is also available in a GitHub repository: MargauxLefebvre/ViroSeek

