## Supplementary data for "ViroSeek: a viral detection pipeline for second-generation sequencing"

### Supplementary information 1: Libraries preparation for MixA to MixC sample based on “SureSelectXT target Enrichment System for Illumina Paired-End Multiplexed Sequencing”.

### 1. Retro-transcription

Retrotranscription was performed using *SuperScript III First-Strand Synthesis System for RT-PCR* (Invitrogen®)

#### a. Mix 1: Denaturation

| Reagent | Volume for 1 reaction (µL) |
| --- | --- |
| RNA | 8 |
| dNTPs | 1 |
| Random primers | 1 |
| Total | 10 |

- Dispense 2µL per reaction and add 8µL of ARN

#### b. Incubation

| 65°C | 5min |
| --- | --- |
| 4°C | +∞ |

- Transfer to ice

#### c. Mix 2: cDNA synthesis

| Reagent | Volume for 1 reaction (µL) |
| --- | --- |
| Buffer 10X | 2 |
| MgCl2 25nM | 4 |
| DTT 0.1M | 2 |
| RNase OUT 40 U/µL | 1 |
| Superscript III RT | 1 |
| Total | 10 |

- add 10µL of mix2 per previous reaction

#### d. Incubation

| 25°C | 10min |
| --- | --- |
| 50°C | 50min |
| 85°C | 5min |
| 4°C | +∞ |

### 2. Second strand synthesis

Second strand synthesis was performed using NEBNext® Ultra™II Non-directional RNA second strand synthesis module (E6111) protocol from NEB

#### a. Mix

| Reagent | Volume for 1 reaction (µL) |
| --- | --- |
| cDNA | 20 |
| NEBNext Second Strand Synthesis Reaction Buffer | 8 |
| NEBNext Second Strand Synthesis Enzyme Mix | 4 |
| Nuclease free water | 48 |
| Total | 80 |

- Mix by pipetting 10 times back and forth
- Dispense 60µL per reaction and add 20µL of cDNA

#### b. Incubation

| 16°C | 1h |
| --- | --- |
| 4°C | +∞ |

- Do not let thermocycler lid heat up more than 40°C

### 3. Purification N°1

Purification has been performed using *AMPure XP* beads from Beckman Coulter®

- Bring beads to room temperature 30 min before use
- Re-suspend dsDNA with 1. 8X beads, i.e. 144µL, by pipetting 10 times back and forth
- Incubate 5 min at RT then place on the magnetic rack
- Carefully remove the supernatant and add 200µL of 80% ethanol, without taking up the beads
- Incubate 30s and repeat the wash again
- Remove the supernatant and leave the tube open on the rack for 5 min
- If there is no more liquid, but no cracking of the beads, add 31µL of water
- Incubate 2 min, reposition on the magnetic rack for 2-3 min and transfer the supernatant to a tube 1. 5mL

### 4. Enzymatic fragmentation

Enzymatic fragmentation has been performed using *FS DNA library prep Kit* (E7805, E6177) protocol with inputs ≥100 ng from NEB

#### a. Fragmentation mix

| Reagent | Volume for 1 reaction (µL) |
| --- | --- |
| DNA | 26 |
| NEBNext Ultra II FS Reaction buffer | 7 |
| NEBNext Ultra II FS Enzyme Mix | 2 |
| Total | 35 |

#### b. Incubation

| 37°C | 10min |
| --- | --- |
| 65°C | 30min |
| 4°C | +∞ |

### 5. Purification N°2

The purification is the same as in step 3, with the bead ratio remaining 1.8×. The elution is performed in 15 µL.

### 6. Ligation

Ligation has been performed using *SureSelect XT Library Prep kit ILM*

#### a. Mix 1: SureSelect Adaptor Oligo mix 1/10

| Reagent | Volume for 1 reaction (µL) |
| --- | --- |
| Nuclease free water | 1 |
| SureSelect Adaptor Oligo | 9 |
| Total | 10 |

#### b. Mix 2: Ligation Master Mix

| Reagent | Volume for 1 reaction (µL) |
| --- | --- |
| Nuclease free water | 15.5 |
| 5X T4 DNA Ligase Buffer | 10 |
| Mix 1 (SureSelect Adaptor Oligo mix 1/10) | 10 |
| T4 DNA ligase | 1.5 |
| Total | 37 |

- Add 13 of fragmented DNA to the 37µL ligation mix
- Mix by pipetting 10 times back and forth

#### c. Incubation

| 20°C | 15min |
| --- | --- |
| 4°C | +inf. |

### 7. Purification N°3

The purification is the same as in step 3, with the bead ratio remaining 1.8×. 80% ethanol has been replaced by 70% ethanol The elution is performed in 34µL.

### 8. Quantification

Quantification was performed using Qubit dsDNA High Sensibility Assay Kit

| Sample | MixA | MixB | MixC |
| --- | --- | --- | --- |
| Sample concentration (ng/µL) | 1.66 | 1.14 | 1.26 |

### 9. PCR pre capture

#### a. Mix

| Kit | Reagent | Volume for 1 reaction (µL) |
| --- | --- | --- |
|  | Nuclease free water | 6 |
| SureSelect XT Library Prep kit ILM | SureSelect Primer | 1.25 |
| SureSelect Target Enrichment kit IKM Indexing Hyb Module Box 2 | SureSelect ILM Indexing PreCapture PCR Reverse Primer | 1.25 |
| Herculase II Fusion DNA Polymeras kit | 5X Herculase II Reaction Buffer | 10 |
| Herculase II Fusion DNA Polymeras kit | 100mM dNTP Mix | 0.5 |
| Herculase II Fusion DNA Polymeras kit | Herculase II Fusion DNA Polymerase | 1 |
|  | Total | 20 |

- Add 30µL of ligated DNA to the 20µL mix
- Mix by pipetting 10 times back and forth

#### b. Incubation

| 98°C | 2min |  |
| --- | --- | --- |
| **98°C** | **30sec** | **X 14 cycles** |
| **65°C** | **30sec** |  |
| **72°C** | **1min** |  |
| 72°C | 10min |  |
| 4°C | +∞ |  |

### 10. Purification N°4

The purification is the same as in step 7, with the bead ratio remaining 1.8×. The elution is performed in 30 µL.

### 11. Quantification

Quantification was performed using Qubit dsDNA High Sensibility Assay Kit

| DJ: | MixA | MixB | MixC |
| --- | --- | --- | --- |
| Sample concentration (ng/µL) | 48.0 | 54.0 | 54.0 |

### 12. Hybridization and capture

#### a. Mix 1: Hybridization Buffer

| Kit | Reagent | Volume for 1 reaction (µL) |
| --- | --- | --- |
| SuperSelect Target Enrichment Box 1 | SureSelect Hyb 1 | 6.63 |
| SuperSelect Target Enrichment Box 1 | SureSelect Hyb 2 | 0.27 |
| SureSelect Target Enrichment kit ILM Indexing Hyb Module Box 2 | SureSelect Hyb 3 | 2.65 |
| SuperSelect Target Enrichment Box 1 | SureSelect Hyb 4 | 3.45 |
|  | Total | 13 |

- Keep at room temperature

#### b. Mix 2: SureSelect Block Mix

SureSelect Block mix was prepared with SureSelect Target Enrichment kit ILM Indexing Hyb Module Box 2

| Reagent | Volume for 1 reaction (µL) |
| --- | --- |
| SureSelect Indexing Block 1 | 2.5 |
| SureSelect Block 2 | 2.5 |
| SureSelect ILM Indexing Block 3 | 0.6 |
| Total | 5.6 |

- Keep on ice until used

#### c. Mix 3: RNase Block dilution 1/10 Mix

RNase block dilution mix was prepared with SureSelect Target Enrichment KIT ILM Indexing Hyb Module Box 2

| Reagent | Volume for 1 reaction (µL) |
| --- | --- |
| Nuclease free water | 4.5 |
| RNase Block | 0.5 |
| Total | 5 |

- Keep on ice until used
- Add mix 2 ( SureSelect Block mix) to 4.4 ligated DNA

#### d. Incubation

| 95°C | 5min |
| --- | --- |
| 65°C | 5min minimum |

- Pause the program

#### e. Mix 4 final: Capture library and hybridization mix

| Reagent | Volume for 1 reaction (µL) |
| --- | --- |
| Mix1 (Hybridization buffer) | 13 |
| Mix 3 (RNase Block 1/10 mix) | 5 |
| Capture Library (diluted to 1/5) | 1 |
| Total | 19 |

- Keep at room temperature until used (used as fast as possible)
- Add 19µL of mix4 directly in the sample at 65°C, mix by pipetting 10 times back and forth

#### f. Incubation

| 65°C | 16h (overnight) |
| --- | --- |

### 13. Purification post-capture

Purification was performed using Dynabeads MyOne Streptavidin T1 (Invitrogen®)

Incubate 1.5mL of wash buffer 2 at 65°C for future used

#### a. Preparation of the beads

- Vortex the beads thoroughly
- In 1.5mL tube, add 50µL of beads
- Add 200µL of SureSelect Binding Buffer, pipette thoroughly
- Place on the magnetic stand and remove the supernatant
- Repeat this wash step twice
- Resuspend the beads in 200µL of SureSelect Binding Buffer

#### b. Capture of Hybridized DNA

- Add 29µL of hybridization mix (keeping it at a maximum of 65°C until collection) to the 200µL of washed streptavidin beads.
- Mix slowly back and forth until the beads are completely resuspended.
- Place the tube in a thermocycler for 30 min at 1400 rpm at RT.
- After a brief spin, place on a magnetic stand and remove the supernatant.
- Resuspend the beads with 200µL of SureSelect Wash Buffer 1.
- Incubate for 5 min at RT, place back on the magnetic stand and remove the supernatant.

#### c. Washing the beads (maintain a temperature of 65°C)

- Use Wash Buffer 2 preheated to 65°C
- Thoroughly resuspend the beads in 200µL of Wash Buffer 2
- Incubate for 5 minutes at 65°C, return to a magnetic support, and remove the supernatant
- Repeat this wash for a total of 6 washes
- When there is no longer any trace of buffer, resuspend the beads in 30µL of water
- Keep on ice

### 14. PCR post Capture

#### a. Post capture PCR reaction mix

| Kit | Reagent | Volume for 1 reaction (µL) |
| --- | --- | --- |
|  | Nuclease free water | 18.5 |
| Herculase II Fusion DNA Polymeras kit | 5X Herculase II Reaction Buffer | 10 |
| Herculase II Fusion DNA Polymeras kit | Herculase II Fusion DNA Polymerase | 1 |
| Herculase II Fusion DNA Polymeras kit | 100mM dNTP Mix | 0.5 |
| SureSelect Target Enrichment kit ILM Indexing Hyb Module Box 2 | SureSelect ILM Indexing Post-Capture Forward PCR Primer | 1 |
|  | Total | 31 |

- Add 31µL of post capture mix in a new tube
- Add 5µL of indexing primer for each primer
- Add 14µL of DNA (with beads) and pipet all the volume (final volume= 50µL)

#### b. Incubation

| 98°C | 2min |  |
| --- | --- | --- |
| **98°C** | **30sec** | **X 25 cycles** |
| **57°C** | **30sec** |  |
| **72°C** | **1min** |  |
| 72°C | 10min |  |
| 4°C | +∞ |  |

### 15. Purification N°5

16. The purification is the same as in step 3, with the bead ratio remaining 1.8×. The elution is performed in 30 µL.

### Supplementary information 2: Library preparation for AltMix sample based on « Twist Library Preparation Kit for ssRNA Virus Detection (version august 2020) and Twist Target Enrichment Protocol (version November 2021) from Twist Bioscience », with modifications.

### 1. Retro-transcription

Retrotranscription was performed using *SuperScript III First-Strand Synthesis System for RT-PCR* (Invitrogen®)

#### a. Mix 1: Denaturation

| Reagent | Volume for 1 reaction (µL) |
| --- | --- |
| RNA | 8 |
| dNTPs | 1 |
| Random primers | 1 |
| Total | 10 |

- Dispense 2µL per reaction and add 8µL of ARN

#### b. Incubation

| 65°C | 5min |
| --- | --- |
| 4°C | +∞ |

- Transfer to ice

#### c. Mix 2: cDNA synthesis

| Reagent | Volume for 1 reaction (µL) |
| --- | --- |
| Buffer 10X | 2 |
| MgCl2 25nM | 4 |
| DTT 0.1M | 2 |
| RNase OUT 40 U/µL | 1 |
| Superscript III RT | 1 |
| Total | 10 |

- add 10µL of mix2 per previous reaction

#### d. Incubation

| 25°C | 10min |
| --- | --- |
| 50°C | 50min |
| 85°C | 5min |
| 4°C | +∞ |

### 2. Second strand synthesis

Second strand synthesis was performed using NEBNext® Ultra™II Non-directional RNA second strand synthesis module (E6111) protocol from NEB

#### a. Mix

| Reagent | Volume for 1 reaction (µL) |
| --- | --- |
| cDNA | 20 |
| NEBNext Second Strand Synthesis Reaction Buffer | 8 |
| NEBNext Second Strand Synthesis Enzyme Mix | 4 |
| Water | 48 |
| Total | 80 |

- Mix by pipetting 10 times back and forth
- Dispense 60µL per reaction and add 20µL of cDNA

#### b. Incubation

| 16°C | 1h |
| --- | --- |
| 4°C | +∞ |

- Do not let thermocycler lid heat up more than 40°C

### 3. Purification N°1

Purification has been performed using *AMPure XP* beads from Beckman Coulter®

- Bring beads to room temperature 30 min before use
- Re-suspend dsDNA with 1. 2X beads, i.e. 96µL, by pipetting 10 times back and forth
- Incubate 5 min at RT then place on the magnetic rack
- Carefully remove the supernatant and add 200µL of 80% ethanol, without taking up the beads
- Incubate 30s and repeat the wash again
- Remove the supernatant and leave the tube open on the rack for 5 min
- If there is no more liquid, but no cracking of the beads, add 27µL of water
- Incubate 2 min, reposition on the magnetic rack for 2-3 min and transfer 25µL of the supernatant to a tube 1. 5mL

### 4. Enzymatic fragmentation

Enzymatic fragmentation has been performed using *Twist Library preparation EF Kit* (Part 100572 – Twist Bioscience®).

#### a. Mix

| Reagent | Volume for 1 reaction (µL) |
| --- | --- |
| Nuclease free water | 10 |
| 10X fragment buffer | 5 |
| 5X fragmentation enzyme | 10 |
| Total | 25 |

- Add 25µL of mix to the 25µL of dsDNA
- Mix by pipetting 10 times back and forth
- Place in the thermocycler

#### b. Incubation

| 32°C | 14min |
| --- | --- |
| 65°C | 30min |
| 4°C | +∞ |

- The lid heats to 70°C

### 5. Ligation

Ligation has been performed using *Twist Library preparation EF kit 1* (Part 100572 - Twist Bioscience) and *Twist universal adapter system* (Part 101308 - Twist Bioscience®).

- Prepare 1mL of 80% ethanol per sample for the next purification steps.
- Prepare the thermocycler at 20°C, without heating the lid
- ADD 2.5µL of Twist universal adapters to the dA-tailed DNA and keep on ice.

#### a. Mix

| Reagent | Volume for 1 reaction (µL) |
| --- | --- |
| Nuclease free water | 17.5 |
| DNA ligation buffer | 20 |
| DNA ligation mix | 10 |
| Total | 47.5 |

- Add 47.5µL of mix to the 52.5µL of dsDNA+adapters and mix well by pipetting..

#### b. Incubation

| 20°C | +∞ (preheat) |
| --- | --- |
| 20°C | 15min |

### 6. Purification N°2

The purification is the same as in step 3, with the bead ratio remaining 0.8×. The elution is performed in 15 µL.

### 7. PCR with unique dual index (UDI)

PCR was performed using KAPA Hifi HotStart Readymix (KK2602, Kapa Biosystems) and UDI primers from Plate A of the Twist universal adapter system (Part 101308 - Twist Bioscience).

- Add 10µL of primers and mix by pipetting
- Add 25µL of KAPA Hifi HotStart Readymix and mix by pipetting

#### a. Incubation

| 98°C | 45sec |  |
| --- | --- | --- |
| **98°C** | **15sec** | **X 14 cycles** |
| **60°C** | **30sec** |  |
| **72°C** | **30sec** |  |
| 72°C | 1min |  |
| 4°C | +∞ |  |

- The lid heats to 105°C

### 8. Purification N°3

The purification is the same as in step 3, with the bead ratio remaining 0.8×. The elution is performed in 22 µL.

### 9. Preparation of libraries for capture (sample pool of 8)

During this step, an appropriate amount of each indexed library is aliquoted to prepare for the hybridization reaction. Both single- and multiplexed libraries (up to 8 indexed libraries per pool) can be used. However, the amount of each indexed library to use depends on the number of samples per pool (Please refer to the table below).

| Number of samples per pool | Quantity of each sample per pool | Total per pool |
| --- | --- | --- |
| 1 | 500ng | 500ng |
| 2 | 500ng | 1000ng |
| 3 | 500ng | 1500ng |
| 4 | 375ng | 1500ng |
| 8 | 187.5ng | 1500ng |

*Table 1 Quantity of each library for different pool sizes according to Twist Bioscience.*

If the amount of indexed library is insufficient, a small amount can be used but with the risk of decreased library complexity. More than 1,500 ng of DNA library can be used. However, the enrichment performance might be reduced if using over 4 μg of DNA input.

Thaw on ice:

- Indexed library pools
- Pan-Viral probes
- Hybridization Mix
- Hybridization Enhancer
- Set a heating block to 65 °C.
- Set a thermal cycler to 65 °C with the heated lid at 85°C.
- Set a thermal cycler to 95 °C with the heated lid at 105°C.
- Heat the Hybridization Mix at 65 °C for 15 minutes on a heating block, then cool to room temperature for 5 minutes.
- Dilute 2 μl of pan-viral probes into 18 μl of nuclease-free water (0.1x)
- Prepare “probe solution”, then mix well by flicking the tube

| Reagent | Volume for 1 reactions (µL) |
| --- | --- |
| Nuclease free water | 6 |
| Hybridization mix | 20 |
| Pan-viral probe (dilute 1:10) | 2 |
| Total | 28 |

- Heat the probe solution to 95 °C for 2 minutes with the thermal cycler lid at 105 °C, then immediately cool on ice for 5 minutes
- While the probe solution is cooling on ice, heat the resuspended libraries at 95 °C for 5 minutes in a thermal cycler with the lid at 105 °C, then equilibrate both probe solution and indexed library to room temperature for 5 minutes
- Add 28 μl of the probe solution to each resuspended indexed library pool, vortex, and pulse-spin.
- Add 30 μl of Hybridization Enhancer directly on top of the entire capture reaction for each library, pulse-spin, and seal the tubes properly.
- Incubate the hybridization reaction at 65 °C for 16 hours with the heated lid at 85 °C.

Important: Seal the tubes tightly to prevent excess evaporation.

### 10. Capture (suite)

30min before the end of the incubation, prepare:

- 200µL de « Wash buffer 1 » at RT
- 700µL de « Wash buffer 2 » at 65°C
- Put at room temperature *streptavidin binding beads* and *DNA purification beads*
- Vortex well the streptavidin binding beads until well-mixed, and add 100 μL to a new 1.5-ml tube for each hybridized reaction pool.
- Add 200 μL of Binding Buffer to each tube, and mix well by pipetting.
- Place the tubes on the magnetic stand for 1 minute, then remove and discard the clear supernatant.
- Remove the tubes from the magnetic stand, and repeat the washing process (steps 3.2. and 3.3.) two more times for a total of 3 times.
- Finally, add 200 μL of Binding Buffer, resuspend the beads, and vortex until homogenized.
- After the hybridization process is completed, open the thermal cycler lid, and directly transfer the volume of each hybridization reaction into a corresponding tube of washed streptavidin binding beads, then mix by pipetting and flicking the tube.

**Important:** Rapid transfer from the thermal cycler is critical to minimize off-target binding. Therefore, the hybridization reaction tubes can be kept on the thermal cycler while transferring their contents to their corresponding washed streptavidin binding bead tubes.

- Transfer the tubes to a shaker or a rotator, and mix the hybridization reaction with streptavidin binding beads for 30 minutes at room temperature, and with a speed of ~1,500 rpm.

**Note:** Do not vortex

- Pulse-spin, and place the tubes on the magnetic stand for 1 minute.
- Remove and discard the clear supernatant with the Hybridization Enhancer without disturbing the bead pellet.

**Note:** The Hybridization Enhancer might still be present after the removal of the supernatant. However, it will not affect the final result.

- Remove the tubes from the magnetic stand, add 200 μL of Wash Buffer 1, mix well by pipetting, and pulse-spin
- Transfer the entire volume of each sample from step 3.10 into a new 1.5-ml microcentrifuge tube.
- Place the tubes on a magnetic stand for 1 minute

**Important:** This step reduces background caused by non-specific banding to the surface of the tube.

- Remove and discard the supernatant
- Remove the tubes from the magnetic stand, add 200 μL of the pre-heated Wash Buffer 2, mix well by pipetting, and pulse-spin
- Incubate the tubes for 10 minutes at 65 °C
- Place them on the magnetic stand for 1 minute
- Remove and discard the clear supernatant
- Repeat the washing process two more times for a total of three washes. Use a 10-μL pipette, and withdraw all the remaining supernatant
- Remove the tubes from the magnetic stand, and add 45 μL of nuclease-free water. Mix well by pipetting, and transfer to ice. The solution hereafter is referred to as the Streptavidin Binding Slurry.

**Note:** From the hybridization reaction to the binding of hybridized targets to streptavidin beads, some modifications were brought to optimize the target enrichment process.

### 11. PCR post capture

Thaw on ice and pulse-vortex the following reagents:

- KAPA HiFi HotStart ReadyMix
- Amplification primers ILMN (Twist Bioscience)
- Prepare 500 μL of 80% ethanol for each Streptavidin Binding Slurry.
- Mix the Streptavidin Binding Slurry well, transfer 22.5 μL in a PCR strip-tube and keep on ice.

**Note:** The remaining Streptavidin Binding Slurry can be stored at -20 °C for future use.

- Prepare a PCR master mix with the following reagents

#### a. Mix

| Reagent | Volume for 1 reaction (µL) |
| --- | --- |
| Nuclease free water | 10 |
| 10X fragment buffer | 5 |

- add 27.5 μL to each Streptavidin Binding Slurry.
- Mix well by pipetting, pulse spin, and run the program below

#### b. Incubation

| 98°C | 45sec |  |
| --- | --- | --- |
| **98°C** | **15sec** | **X20 cycles** |
| **60°C** | **30sec** |  |
| **72°C** | **30sec** |  |
| 72°C | 1min |  |
| 4°C | +∞ |  |

- The lid heats to 105°C

### 12. Purification N°4

Purification is carried out using the ‘DNA purification beads’ in the Twist Bioscience kit.

- Bring the beads to room temperature 30 min before use
- Vortex the beads well
- Transfer the samples to a 1.5mL tube
- Add 1X beads, i.e. 50µL, mixing well by vortexing
- Incubate 5 min at RT, spin down then put on the magnetic rack until a clear supernatant is obtained (1 min)
- Gently remove the SN and put 200µL of 80% ethanol, without taking up the beads
- Incubate for 30s and repeat the wash once more
- Remove the SN (including the last microlitres at P10) and leave the tube open on the rack for 5-10 min without cracking the beads
- If there is no more liquid, but no cracking of the beads, remove the pellet (from the magnetic holder) with 32µL of water
- Incubate for 2 min, reposition on the magnetic holder for 3 min and transfer 30µL to a new 1.5mL tube.

Supplementary figure 1 : Viral relative abundance from targeted families with ViroSeek and an identity threshold at 90%

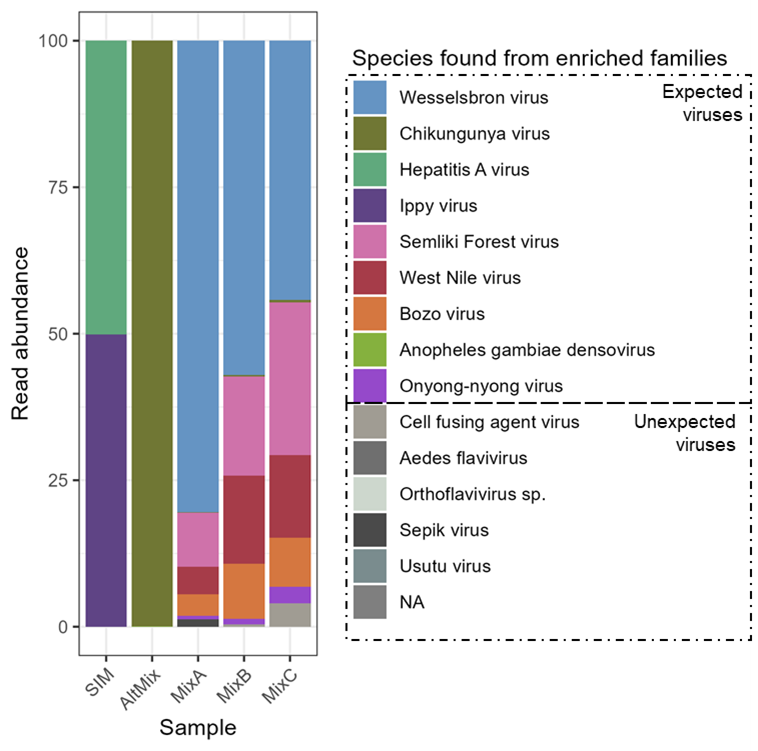

Supplementary table 1: Tools and software versions implemented in the ViroSeek pipeline (v.0.0.2).

| **Tools** | **Version** |
| --- | --- |
| *bbmap* | 39.18 |
| *bioawk* | 1.0 |
| *diamond* | 2.1.13 |
| *fastp* | 0.23.4 |
| *fastqc* | 0.12.1 |
| *minimap2* | 2.29 |
| *samtools* | 1.21 |
| *seqkit* | 2.13.0 |
| *spades* | 4.2.0 |
| *taxonkit* | 0.20.0 |
| *trimgalore* | 0.6.10 |
| *wget* | 1.25 |

Supplementary table 2: Viral species and strains covered in the viral probes panel.

Supplementary table 3: The abundance distribution of the simulated metagenomic dataset generated with *CAMISIM*.

| **Simulation** | **Genomes** | **Distribution** |
| --- | --- | --- |
| 0 | Ippy virus segment S | 0.72 |
|  | Ippy virus segment L | 0.01 |
|  | Hepatitis A virus | 0.21 |
|  | *Helicobacter hepaticus* | 0.06 |
| 1 | Ippy virus segment S | 0.025 |
|  | Ippy virus segment L | 0.935 |
|  | Hepatitis A virus | 0.03 |
|  | *Helicobacter hepaticus* | 0.01 |
| 2 | Ippy virus segment S | 0.005 |
|  | Ippy virus segment L | 0.015 |
|  | Hepatitis A virus | 0.940 |
|  | *Helicobacter hepaticus* | 0.06 |
| 3 | Ippy virus segment S | 0.6 |
|  | Ippy virus segment L | 0.03 |
|  | Hepatitis A virus | 0.04 |
|  | *Helicobacter hepaticus* | 0.33 |
| 4 | Ippy virus segment S | 0.15 |
|  | Ippy virus segment L | 0.12 |
|  | Hepatitis A virus | 0.15 |
|  | *Helicobacter hepaticus* | 0.65 |
| 5 | Ippy virus segment S | 0.08 |
|  | Ippy virus segment L | 0.12 |
|  | Hepatitis A virus | 0.15 |
|  | *Helicobacter hepaticus* | 0.65 |
| 6 | Ippy virus segment S | 0.99 |
|  | Ippy virus segment L | 0.005 |
|  | Hepatitis A virus | 0.003 |
|  | *Helicobacter hepaticus* | 0.002 |
| 7 | Ippy virus segment S | 0.01 |
|  | Ippy virus segment L | 0.03 |
|  | Hepatitis A virus | 0.05 |
|  | *Helicobacter hepaticus* | 0.91 |
| 8 | Ippy virus segment S | 0.07 |
|  | Ippy virus segment L | 0.002 |
|  | Hepatitis A virus | 0.003 |
|  | *Helicobacter hepaticus* | 0.925 |
| 9 | Ippy virus segment S | 0.308 |
|  | Ippy virus segment L | 0.047 |
|  | Hepatitis A virus | 0.254 |
|  | *Helicobacter hepaticus* | 0.391 |
